# High glucose enhances antigen-independent CTL killing via TRAIL

**DOI:** 10.1101/2021.08.04.455060

**Authors:** Wenjuan Yang, Andreas Denger, Caroline Diener, Frederic Küppers, Leticia Soriano-Baguet, Gertrud Schäfer, Archana K. Yanamandra, Renping Zhao, Arne Knörck, Eva C. Schwarz, Martin Hart, Frank Lammert, Leticia Prates Roma, Dirk Brenner, Grigorios Christidis, Volkhard Helms, Eckart Meese, Markus Hoth, Bin Qu

## Abstract

Cytotoxic T lymphocytes (CTLs) are involved in development of diabetes. However, the impact of excessive glucose on CTL-mediated antigen-independent killing remains elusive. Here, we report that TNF-related apoptosis inducing ligand (TRAIL) is substantially up- regulated in CTLs in environments with high glucose (HG) both *in vitro* and *in vivo*. The PI3K- Akt-NFκB axis and non-mitochondrial reactive oxygen species are essential in HG-induced TRAIL upregulation in CTLs. TRAIL^high^ CTLs induce apoptosis of pancreatic beta cell line 1.4E7. Metformin and Vitamin D synergistically reduce HG-enhanced expression of TRAIL in CTLs and coherently protect 1.4E7 cells from TRAIL-mediated apoptosis. Notably, in patients with diabetes, correlation between Vitamin D concentrations in plasma and glucose levels is linked to HG-enhanced TRAIL expression on CTLs. Microarray data reveal that OXCT2, an important enzyme in ketone body catabolism, is a promising target in response to vitamin D. Our work not only reveals a novel mechanism of CTL involvement in progression of diabetes, but also establishes CTLs as a target for combined metformin and vitamin D therapy to protect pancreatic beta cells of diabetic patients.

## Introduction

Cytotoxic T lymphocytes (CTLs) are the key players in the adaptive immune system to destroy pathogen-infected or tumorigenic cells ^1^. Usually, CTLs employ two major killing mechanisms: lytic granules (LGs) and Fas/Fas ligand (FasL) pathway. LGs contain cytotoxic proteins, such as the pore-forming protein perforin and serine proteases granzymes. Upon target recognition, LGs are reoriented towards the CTL-target contact site, termed the immunological synapse (IS), and their cytotoxic content will be specifically released into the cleft, resulting in direct lysis or apoptosis of target cells ^2^. FasL, a member of tumor necrosis factor (TNF) family, is expressed on the surface of CTLs. Engagement of FasL with its Fas receptor, which spans on the target cell surface, initiates apoptotic cascades in target cells ^2, 3^. Emerging evidence shows that the effector functions of CTLs are highly dependent on their metabolic status ^4^. Glucose metabolism and the proper access to glucose are essential to maintain CTL effector functions, especially their killing function ^5^. It is reported that with excessive glucose, calcium influx elicited upon conjugation with target cells is reduced ^6^ and CTL killing efficiency is elevated partially regulated by Ca^2+^ without affecting lytic granule pathway and FasL expression ^7^

Elevated level of blood glucose is a typical symptom for diabetes, a metabolic disease. Diabetes affects 422 million people globally and has become the seventh leading cause of death worldwide ^8^. Diabetes is mainly categorized into two groups, type 1 diabetes and type 2 diabetes. Type 1 diabetes has been identified as an autoimmune disease, for which CTLs play an important role in destroying the insulin-producing pancreatic beta-cells in an antigen specific manner ^9^. For Type 2 diabetes, CTLs, along with other factors, are reported to be associated with its initiation and progression ^10^.

TNF-related apoptosis-inducing ligand (TRAIL), similar as FasL, also belongs to TNF super family. A rich body of studies suggests that TRAIL and/or TRAIL receptors are correlated with progression of diabetes and diabetes-related complications ^11, 12^. Interestingly, TRAIL is used by CTLs to clear viral-infected cells and down-size the effector population to terminate immune responses in an antigen-independent manner ^13^. TRAIL and TRAIL receptors are linked to both type 1 and type 2 diabetes ^14, 15^. Knock-out of TRAIL could protect the high-fat diet-fed hosts from developing diabetes ^16^. In humans, four TRAIL receptors are expressed, TRAIL-R1, -R2, -R3 and -R4 ^17^. The former two (TRAIL-R1 and -R2) are activating receptors, which contain functional death domains to trigger the caspase-8-dependent apoptotic pathway. TRAIL-R3 and -R4 function as decoy receptors, lacking the capability to initiate the death signaling pathway. TRAIL-R1 and/or -R2 are expressed in many normal tissues including pancreas and especially pancreatic beta cells ^18, 19^.

In this work, we show that TRAIL is substantially up-regulated in high glucose-cultured CTLs and in CTLs from diabetic patients or diabetic mouse models. Both the PI3K-Akt-NFκB axis and reactive oxygen species (ROS) are involved in regulating this high glucose-enhanced TRAIL expression. Pancreatic beta-cells can be killed by TRAIL^high^ CTLs in a TRAIL-dependent manner. Furthermore, we found that treatment of two drugs, metformin and vitamin D, could individually or synergistically abolish the enhanced TRAIL expression induced by high glucose in CTLs to protect pancreatic beta cells from TRAIL-mediated apoptosis. Our findings suggest that the uncontrolled high level of blood glucose initiated by diabetes could lead to a TRAL-dependent destruction of beta-cells via CTLs, providing a new insight into the progression of diabetes and a possible intervention strategy to protect beta cells in a context of diabetes.

## Materials and Methods

### Antibodies and reagents

All chemicals are from Sigma-Aldrich (highest grade) if not mentioned otherwise. The following antibodies and reagents were purchased from Biolegend: APC/Cy7 anti-human CD3 antibody, BV421 anti-human CD3 antibody, BV421 anti-human CD8 antibody, APC anti-human CD253 (TRAIL) antibody, APC anti-mouse CD253 (TRAIL) Antibody, BV421 anti-mouse CD8a Antibody, PE anti-mouse CD3 Antibody, PerCP anti-human CD25 antibody, APC anti-human CD62L antibody, APC anti-human CD262 (TRAIL-R2), and 7-AAD viability staining solution. The following antibodies are also used: FITC anti-human CD69 (eBiosciences), FITC anti-human CD44 (DAKO), Purified NA/LE mouse anti-human CD253 (BD Biosciences), BV421 mouse anti-human CD263 (TRAIL-R3) (BD Biosciences), Alexa647 mouse anti-human GLUT1 (BD Biosciences), human TRAIL R1/TNFRSF 10A PerCP-conjugated antibody (R&D Systems), and human TRAIL R4/TNFRSF 10D PE-conjugated antibody (R&D Systems). In addition, the following reagents were used: NucView Caspase-3 enzyme substrates (Biotium), Idelalisib (Selleckchem), MK-2206 (Selleckchem), Rapamycin (Selleckchem), MG-132 (Merck), Caffeic acid phenethyl ester (CAPE) (R&D Systems), Mitoquinone (MitoQ) (Biotrend), N-acetyl-L-cysteine (NAC) (Merck), Vitamin 1,25D3 (Merck), Calcipotriol (TOCRIS), DMSO (Merck), Cellular ROS Assay Kit (Abcam), Metformin hydrochloride (Merck), H2O2 (Merck), and Streptozotocin (Merck).

### Cell culture

Peripheral blood mononuclear cells (PBMCs) were obtained from healthy donors as described elsewhere ^20^. Primary human CD8^+^ T cells were negatively isolated from PBMC using Human CD8^+^ T Cell isolation Kits (Miltenyi Biotec). Human CD8^+^ T cells were stimulated with CD3/CD28 activator beads (Thermo Fisher Scientific) and cultured in DMEM medium (Thermo Fisher Scientific) containing normal (5.6 mM) or high glucose (25 mM) for up to three days if not otherwise mentioned. Levels of glucose in medium was examined every day using “Contour Next Sensoren” test strips (SMS Medipool) and consumed glucose was compensated accordingly. If CTLs were cultured longer than three days, human recombinant IL-2 (Miltenyi Biotec) was added to the medium every two days from day 2 on (100 U/ml). All cells were cultured at 37°C with 5% CO2. Human pancreatic beta cell line 1.4E7 was purchased from Merck and cultured in RPMI-1640 medium (Thermo Fisher Scientific) supplemented with 2 mM glutamine, 1% Penicillin-Streptomycin plus 10% FCS (ThermoFisher Scientific).

### Flow cytometry analysis

For cell surface staining, cells were washed twice with PBS/0.5% BSA and stained for 30 minutes at 4℃ in dark using corresponding antibodies mentioned in the figure legends. For intracellular staining, cells were fixed in pre-chilled 4% PFA and permeabilized with 0.1% saponin in PBS containing 5% FCS and 0.5% BSA, followed by the immunostaining as described above. Flow cytometry data were acquired using a FACSVerse flow cytometer (BD Biosciences) and were analyzed with FlowJo v10 (FLOWOJO, LLC).

### Assays for apoptosis and viability

To assess cell apoptosis, 1.4E7 cell were co-cultured with primary human CD8^+^ T cells and incubated at 37°C with 5% CO2. Cells were harvested at various time points as indicated in the text and stained with BV421-CD3, and then incubated with NucView Caspase-3 Substrates at room temperature for 30 minutes, followed with analysis using flow cytometry. To determine the viability of CTLs, cells were stained with 7-AAD.

### Diabetic mouse model

C57BL/6N mice were injected with streptozotocin intraperitoneally for five days consecutively (50 mg/kg per day) and were sacrificed at day 21. Blood samples were taken from the tail vein and glucose level was tested by standard test strips every day after injection. Mice with blood glucose level more than 250 mg/dL one week after the first injection were considered as diabetic.

### Microarray and analysis

For transcriptome analyses, total RNA of CD8^+^ human T cells was extracted by miRNeasy Mini Kit (Qiagen), following the manufacturers’ instructions, and quantified using NanoDrop 2000c Spectrophotometer (Thermo Fisher Scientific). The quality of the RNA samples was assessed by determining the corresponding RIN (RNA integrity number) values. For this purpose, an Agilent 2100 Bioanalyzer instrument was used together with the RNA 6000 Nano assay from Agilent Technologies (Santa Clara). An excellent quality of the analyzed total RNA was verified by RIN values of 10 for all samples. To determine the cellular transcriptomes, 100 ng of the total RNA was analysed by microarray. The microarray analyses were performed as previously described ^21^. Differential expression analysis of the Agilent microarray data was performed with the Linear Models for Microarray Data (limma) R package. First, samples were corrected for background (NormExp) and quantile normalized, respectively. Control probes used for background correction, probes without associated gene symbols, and genes classified as not expressed in at least 22 out of 24 arrays by the Agilent feature extraction software were removed from the dataset. After the filtering step, the dataset contained a total of 22,658 transcripts, corresponding to 15,360 genes. For differential expression analysis, a linear model was fitted to each individual sample. Then, a linear model was calculated between samples in high glucose (HG) or normal glucose (NG) conditions. Differentially expressed genes were identified with a t-test. The associated p-values were calculated for each gene with an empirical Bayes method and adjusted for multiple testing with the Benjamini-Hochberg method. A gene was classified as differentially expressed (DE) for a given contrast if its adjusted p-value was below 0.05. Among the DE genes, an enrichment analysis for gene annotations was performed with the limma package. Also, an enrichment analysis based on protein interaction subnetworks was carried out by the pathfindR package.

### Seahorse assay

Primary human CD8^+^ T cells were stimulated either in normal (5.6 mM) or high glucose (25 mM) for three days. At day 3, CTLs were counted and seeded in 96-well XF Cell Culture Microplate in XF Seahorse DMEM medium at a cell concentration of 3 x 10^5^ cells/well. Following the manufacturer’s instructions (Agilent), the extracellular acidification rate (ECAR) and oxygen consumption rate (OCR) were measured using the XF Glycolytic Stress Test and XF Cell Mitochondrial Stress Test kits, respectively.

### Patient material

Blood samples were collected from patients with diabetes type 1 and 2 (using the current diagnostic criteria from the American Diabetes Association-ADA) and healthy control subjects (all of which had a normal glycated haemoglobin level (HbA1c < 5,7%) at the day of the blood sample collection). Both patients and healthy controls were recruited in the Department of Internal Medicine 2 in the University Medical Center of Homburg, Saarland, Germany. Written informed consent was obtained from all subjects before the blood sampling, strictly following the procedure described in the ethical approval of the ethic committee of the medical association of Saarland (Ethic Vote Nr: Ha 84/19). PBMCs were obtained from the blood samples and were stimulated with CD3/CD28 activator beads (Thermo Fisher Scientific) and cultured in DMEM containing normal (5.6 mM) or high glucose (25 mM) for three days supplemented with recombinant IL-2 (100 U/ml, Miltenyi).

### Intracellular ROS detection

Cellular ROS Assay Kit (Abcam) was used to determine intracellular ROS. Briefly, at 6 hours after activation of CD3/CD28 activator beads, the CTLs were stained with BV421-CD8 for 30 minutes at 4 °C in dark, and then incubated with dichlorofluorescein diacetate (DCFDA, 20 µM) at 37 °C for 30 minutes, followed by analysis using flow cytometry.

### Quantitative RT-PCR

The mRNA expression analysis was carried out as described before ^22^. Briefly, total RNA was isolated from CTLs using TRIzol reagent (ThermoFisher Scientific). Then the isolated RNA was reversely transcribed into complementary DNA (cDNA) and relative gene expression was performed by qRT-PCR using CFX96Real-TimeSystemC1000 Thermal Cycler (Bio-Rad Laboratories). TATA box-binding protein (TBP) was used as the housekeeping gene for the normalization of the target genes. Primer sequences are as follows (forward/reverse): TBP (5’- CGGAGAGTTCTGGGATTGT-3’/5’-GGTTCGTGGCTCTCTTATC-3’). Pre-designed primers were purchased for TRAIL (QT00068957) and Glut1 (QT00079212).

### Statistical analysis

Data are presented as mean ± SEM. GraphPad Prism Software (San Diego, CA, USA) was used for statistical analysis. The differences between two groups were analyzed by the Student’s t-test. For multiple comparisons, two-way ANOVA or one-way ANOVA was performed followed by Bonferroni test.

## Results

### Expression of TRAIL is up-regulated in CTLs by high glucose and TRAIL^high^ CTLs induces destruction of pancreatic beta cells

To test the molecular mechanism how high glucose increases CTL cytotoxicity, we analyzed the TRAIL pathway, since high glucose does not affect lytic granule pathway and FasL expression ^7^. We used negatively isolated primary human CD8^+^ T cells and stimulated the cells with CD3/CD28 antibody-coated beads in presence of normal (5.6 mM, NG) or high glucose (25 mM, HG) for three days (hereafter referred to as NG or HG- CTLs). At mRNA level, TRAIL was substantially up-regulated in HG-CTLs compared to NG-CTLs (Fig. 1a). Concomitantly, at protein level, not only the total expression of TRAIL was considerably enhanced (Fig. 1b), but also the expression of TRAIL on the surface was significantly elevated in HG-CTLs compared to NG-CTLs (Fig. 1c, d). To test the *in vivo* relevance of this finding, we used a streptozotocin-induced diabetic mouse model. Compared to CTLs from the control group, CTLs from diabetic mice exhibited significantly elevated level of TRAIL (Fig. 1e). Furthermore, the level of blood glucose was positively correlated with expression of TRAIL (Fig. 1f). We conclude that TRAIL expression in CTLs is up-regulated by HG and that glucose levels correlate with TRAIL expression *in vivo* in the context of diabetes.

**Figure 1.**
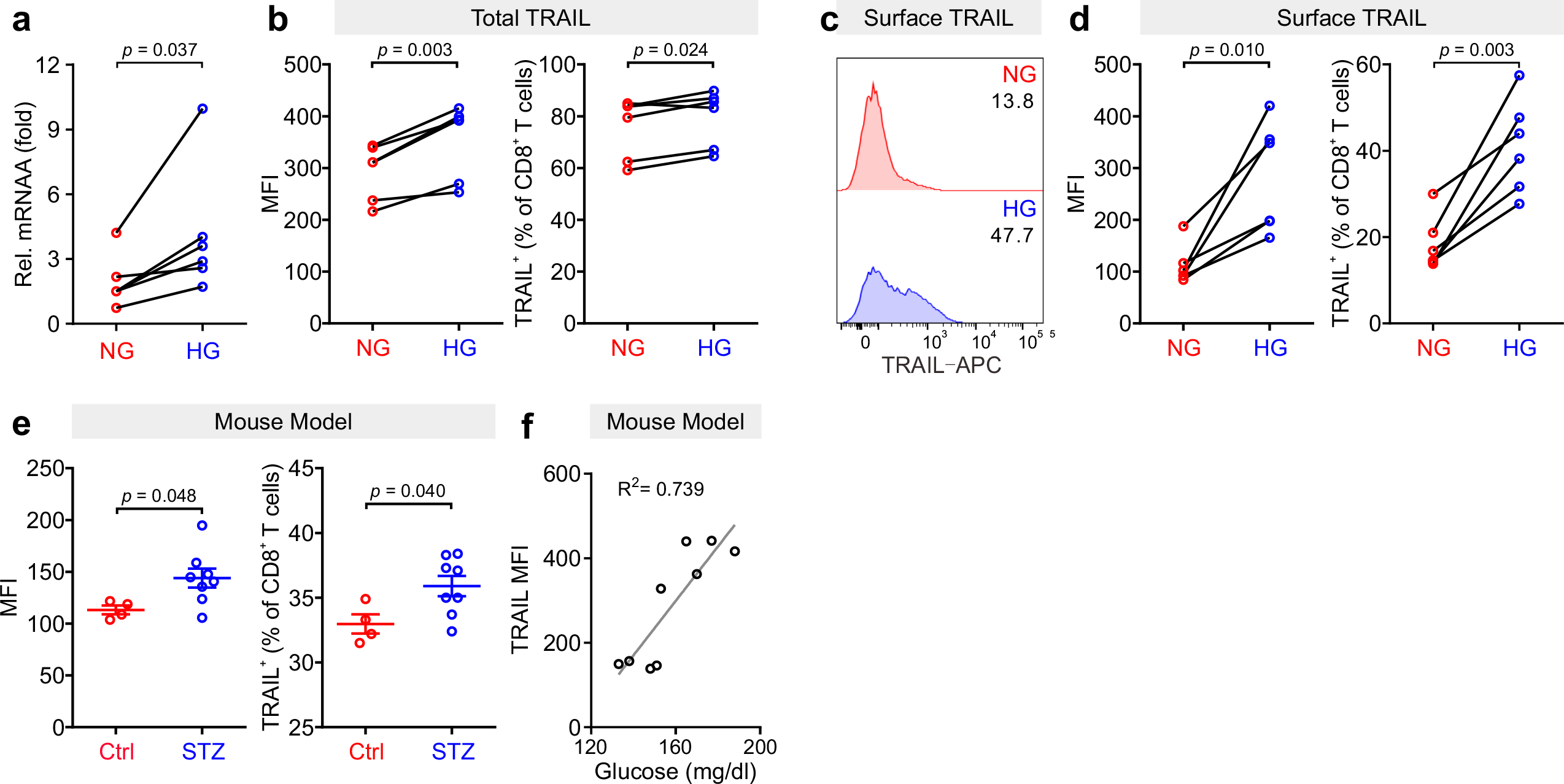
TRAIL is up-regulated in CTLs in environments with high glucose. Primary human CD8^+^ T cells were stimulated with CD3/CD28 beads for 3 days in NG (5.6 mM) or HG (25 mM) medium. (**a**) Relative mRNA expression levels of TRAIL in CTLs were quantified by qRT-PCR (n = 6). (**b-d**) Expression of TRAIL in CTLs was detected by flow cytometry. CTLs were stained with antibody against CD8 and TRAIL following permeabilization (**b**) or without permeabilization (**c**, **d**) for total or surface protein level of TRAIL (n = 6). (**e, f**) TRAIL is up-regulated in diabetic mice. Mouse splenocytes were isolated and stained with PE-mCD3, BV421-mCD8, and APC-mTRAIL for analysis using flow cytometry (Ctrl = 4, STZ mice = 8). Data are represented as Mean ± SEM. Correlation of TRAIL expression in CD8^+^ T cells and blood glucose is shown in **f**. MFI, mean fluorescent intensity. Data were analyzed by two-tailed paired Student’s *t* test **(a**, **b**, and **d**), two-tailed unpaired Student’s *t* test (**e**) or trendline analysis (**f**).

Given the fact that treatment of soluble TRAIL can induce apoptosis of human pancreatic beta cells ^19^, we then examined whether CTLs expressing TRAIL could have a similar effect. We used the human pancreatic beta cell line 1.4E7, which is a hybrid cell line derived from electrofusion of primary human pancreatic islets with a human pancreatic ductal carcinoma cell line PANC-1. We incubated 1.4E7 cells with CTLs and used activity of caspase-3 in 1.4E7 cells as a readout for CTL-induced apoptosis. The results show that HG-cultured CTLs exhibited significantly higher killing capacity compared to their counterparts in NG (Fig. 2a-d; Fig. S1a, b). Of note, apoptosis of beta cells is positively correlated with TRAIL expression (Fig. 2e, f; Fig. S1c). To test a potential causal relation between TRAIL expression and CTL cytotoxicity against beta cells, we blocked TRAIL function with its neutralizing antibody. The analysis of caspase-3 activity shows that TRAIL blockade diminished CTL-mediated killing against 1.4E7 beta cells to a large extent for HG-CTLs (Fig. 2g, h). Next, we examined the expression of TRAIL receptors on 1.4E7 cell surface. We found that out of four TRAIL receptors, TRAIL-R2 was predominantly expressed (Fig. S1d), which likely mediates TRAIL^high^ CTL-induced apoptosis of 1.4E7 cells. Taken together, our results suggest that HG- CTLs destroy pancreatic beta cells in a TRAIL-mediated manner.

**Figure 2.**
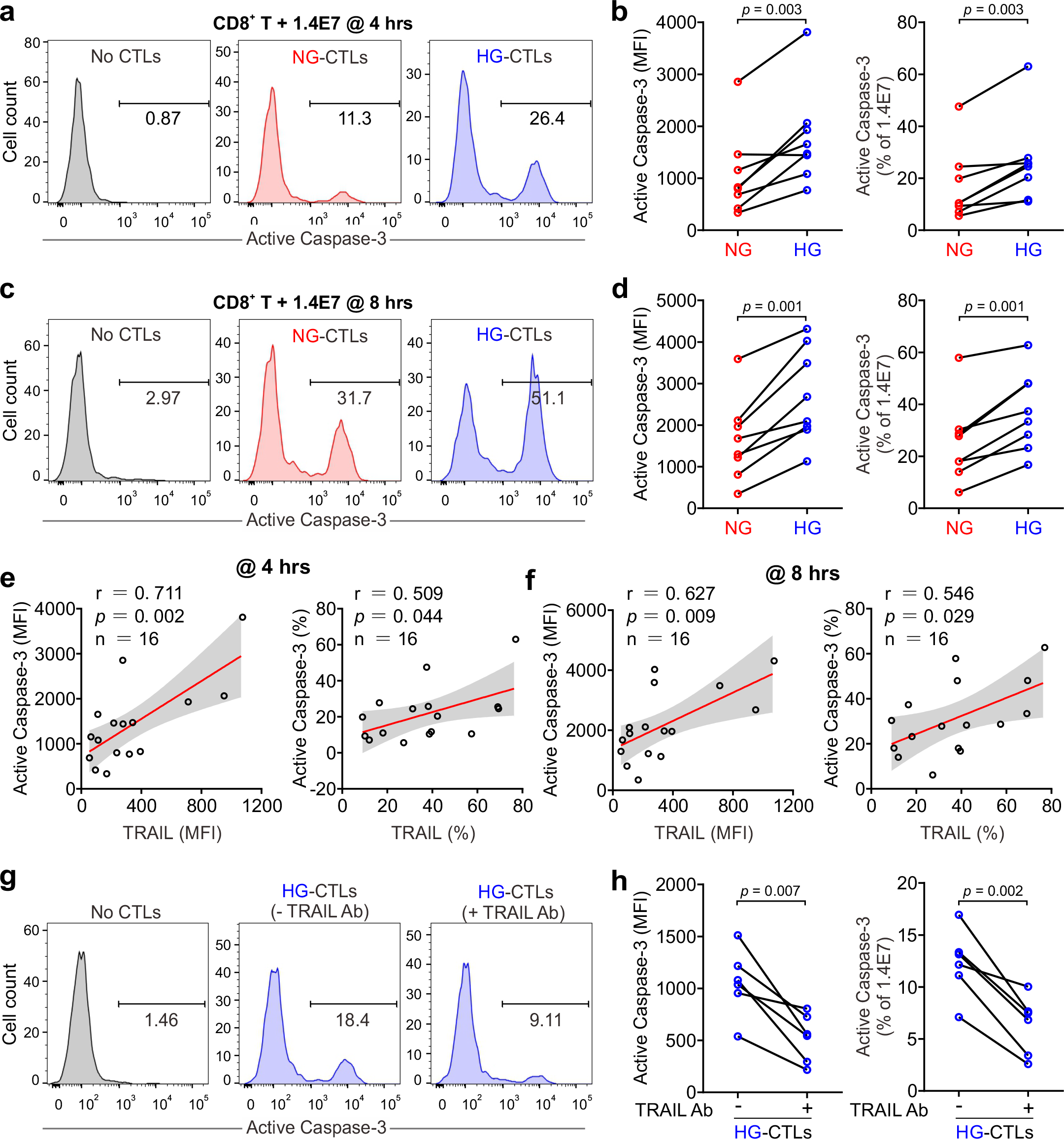
Up-regulation of TRAIL in HG-CTLs enhances apoptosis of pancreatic beta cells. Primary human CD8^+^ T cells were stimulated with CD3/CD28 beads for 3 days in NG (5.6 mM) or HG (25 mM) medium. (**a-d**) Apoptosis of pancreatic beta cells induced by CTLs were measured by staining active Caspase-3 using flow cytometry. Human pancreatic beta cells 1.4E7 were incubated with CTLs with an effector to target (E:T) ratio of 20:1 for 4 hours (**a**, **b**) or 8 hours (**c**, **d**) (n = 8). One representative donor out of eight is shown in **a** and **c**, respectively. (**e, f**) Correlation of expression of TRAIL in CD8^+^ T cells with apoptosis of 1.4E7 cells (Caspase-3 activity) (n = 16). (**g, h**) CD8^+^ T cells-induced beta cell apoptosis is TRAIL dependent. Caspase-3 activity was analyzed in co-culture of 1.4E7 cells and CTLs in the presence or absence of anti-human TRAIL antibody (50 µg/ml) for 4 hours. One representative donor is shown in **g** and the quantification is shown in **h** (n = 6). MFI, mean fluorescent intensity. Data were analyzed by two-tailed paired Student’s *t* test **(b**, **d**, and **h**) or Pearson’s correlation coefficients (**e**, and **f,** and **k**).

### Metabolic processes in CTLs are reprogrammed by high glucose

Glucose metabolism encompasses the intracellular biochemical processes to breakdown and utilize glucose to generate energy, including oxidative phosphorylation and glycolysis. Using the Seahorse assay, we evaluated the oxygen consumption rate (OCR) and the extracellular acidification rate (ECAR). We found that HG-cultured CTLs exhibited significantly elevated OCR and ECAR compared to their counterparts cultured in NG (Fig. 3a-d).

**Figure 3.**
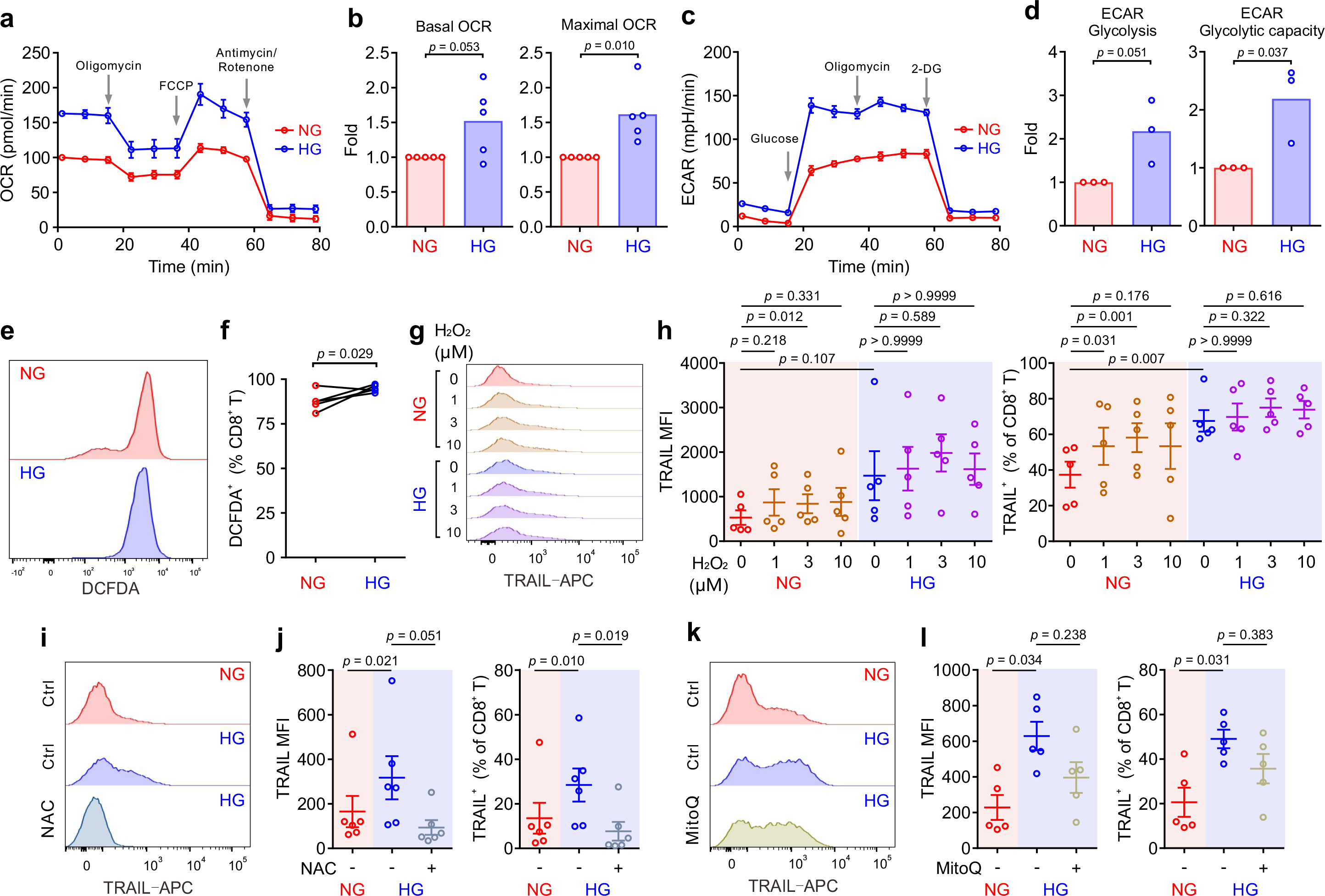
Metabolic processes in CTLs are reprogrammed by high glucose. Primary human CD8^+^ T cells were stimulated with CD3/CD28 beads for 3 days in NG (5.6 mM) or HG (25 mM) medium. (**a-d**) Oxidative phosphorylation (n = 5) and glycolysis (n = 3) of CTLs were determined with seahorse assay. One representative donor for oxidative phosphorylation and glycolysis is shown in **a** and **c**, respectively. (**e, f**) ROS production in CD8^+^ T cells was determined at 6 hours after CD3/CD28 bead stimulation by DCFDA (n = 5). One representative donor is shown in **e**. (**g, h**) H2O2 enhances TRAIL expression in CTLs in NG. CD8^+^ T cells were stimulated with CD3/CD28 beads in presence or absence of H2O2 for 3 days (n = 5). One representative donor is shown in **g**. (**i-l**) Inhibition of ROS production abolishes HG-enhanced TRAIL expression in CTLs. NAC (**i**, **j**, 10 mM, n = 6) or MitoQ (**k**, **l**, 0.4 μM, n = 5) was added during the activation for 3 days. One representative donor for NAC and MitoQ is shown in **i** and **k**, respectively. Results are represented as Mean ± SEM. Data were analyzed by two-tailed unpaired Student’s *t* test (**b** and **d**), two-tailed paired Student’s *t* test (**f**) or one-way ANOVA (**h**, **j** and **l**).

Given the essential role of mTOR in regulating glycolysis in T cells ^23^, we next examined the possible involvement of mTOR in HG-induced metabolic reprogramming in CTLs. To our surprise, no difference was identified between NG- and HG-CTLs regarding mTOR activation (Fig. S2a). Since glucose uptake and transport play an essential role for overall glucose metabolism, we examined the expression of glucose transporter 1 (Glut1) and found that although its expression was down-regulated at mRNA level in HG-CTLs (Fig. S2b), at the protein level it remained unchanged (Fig. S2c, d). In addition, the fraction of Glut1 transported to the plasma membrane was also not altered by HG (Fig. S2e, f). Together, our results suggest that neither mTOR nor glucose transport are likely to be responsible for HG-reprogrammed metabolic processes in CTLs.

To gain deeper insights into the genes affected by HG for metabolic reprogramming, we compared transcriptomes of NG-CTLs and HG-CTLs (Spreadsheet 1). First, we examined the expression of all glucose transporters. We found that three glucose transporters (Glut1/SLC2A1, Glut3/SLC2A3, and Glut14/SLC2A14) were predominantly expressed in CTLs, and moderate downregulation in Glut1 was observed in HG-CTLs compared to their NG counterparts (Fig. S2g), which is in a good agreement with our data from quantitative PCR (Fig. S2b). No further difference in expression of glucose transporters was identified between NG- and HG-CTLs (Fig. S2g), further supporting our conclusion that glucose transport is unlikely to be responsible for HG-reprogrammed metabolic processes in CTLs. With regards to enriched GO terms, the most significant change happened in the cellular metabolism. When comparing HG- to NG-CTL samples, many genes involved in metabolic processes are significantly changed with a total of 295 being significantly up-regulated and 206 genes being significantly down-regulated (Spreadsheet 1). A total of 83 GO annotations describing various metabolic processes are associated with at least one significantly deregulated gene (Spreadsheet 2). Furthermore, 16 genes annotated with the metabolism of ROS were significantly deregulated (Spreadsheet 2).

We then examined whether ROS was involved in HG-enhanced TRAIL expression. We first determined ROS production in CTLs using DCFDA, a fluorescent probe used to detect ROS in living cells ^24^. We found that upon activation, ROS produced in HG- CTLs was higher than that in their NG counterparts (Fig. 3e, f). To explore whether ROS is involved in HG-enhanced TRAIL expression, we first added H2O2, a relative stable form of ROS, during NG- and HG-culture. We found that with additional H2O2, TRAIL expression was elevated in NG-cultured CTLs but not in HG-cultured CTLs (Fig. 3g, h). It implies that the elevation in TRAIL expression by HG is likely via enhanced ROS in HG-CTLs. To verify this hypothesis, we used

ROS scavengers either with a general inhibition effect (N-acetyl-L-cysteine (NAC)) or specifically targeted to mitochondrial ROS (MitoQ). We found that when ROS production in cytosol was removed by NAC the expression of TRAIL was drastically decreased in HG-CTLs to a comparable level as in NG-CTLs (Fig. 3i, j). In contrast, removal of mitochondria-produced ROS by MitoQ in HG-CTLs did not significantly alter TRAIL expression (Fig. 3k, l). Thus, our results suggest that in CTLs, ROS, especially cytosolic ROS, plays a key role in HG-enhanced TRAIL expression.

### HG-enhanced TRAIL expression on CTLs is regulated by the PI3K/Akt/NFκB axis

Transcriptome analysis also revealed a statistically significant up-regulation of 11 genes including TRAIL annotated with the KEGG pathway *apoptosis* in HG samples (Spreadsheet 3). Among the top 10 GO terms affected by HG, TRAIL (synonym name TNFSF10) is linked both to “positive regulation of IκB kinase/NFκB signaling” and to “positive regulation of apoptotic process” (Fig. 4a).

**Figure 4.**
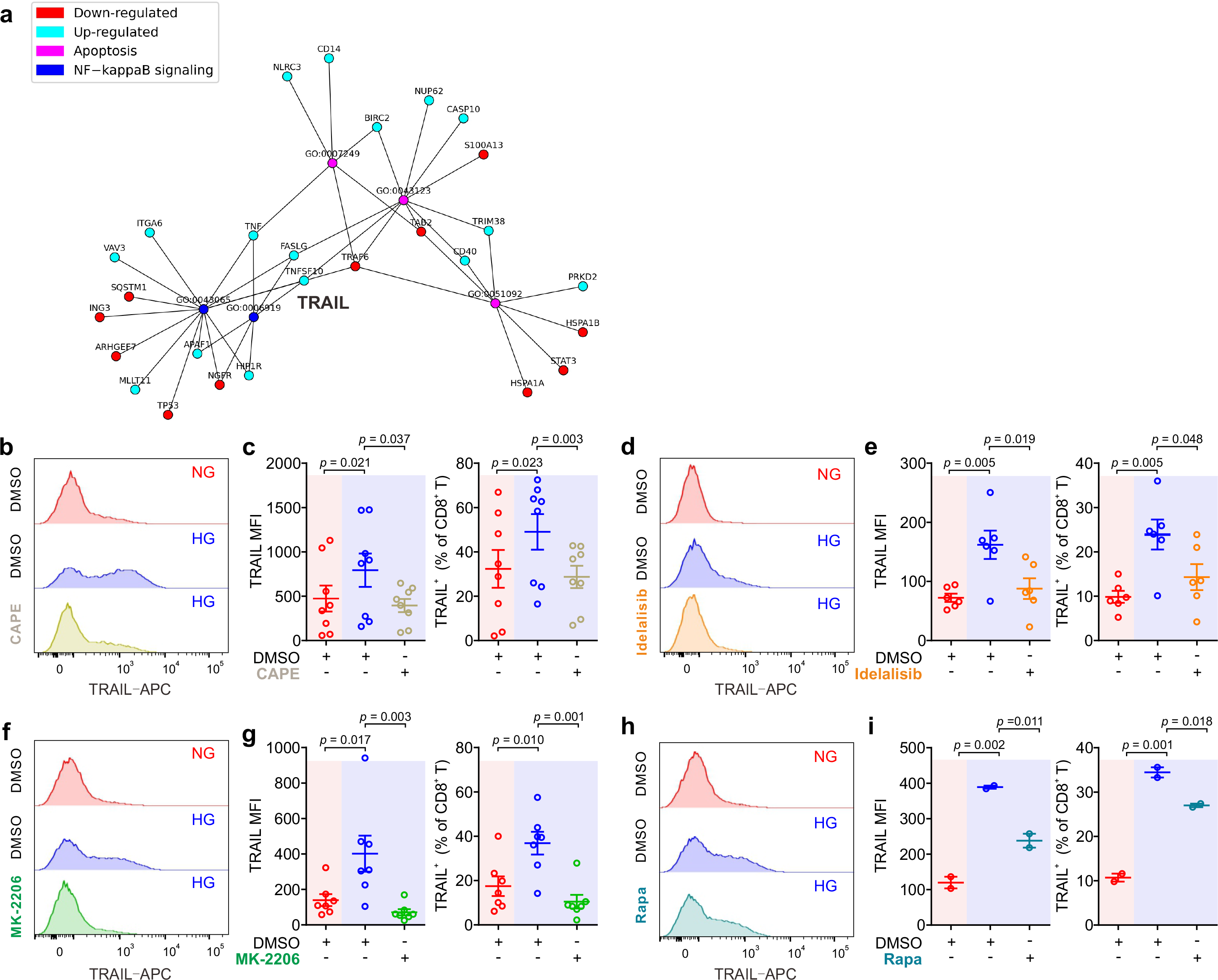
Up-regulation of TRAIL in CTLs is regulated by the PI3K/Akt/NFκB axis. (**a**) Ten most significantly enriched/depleted Gene ontology terms in HG vs NG conditions. Enrichment analysis was performed with the *pathfindR* package, which creates a sub-network of the human protein-protein interaction network that only contains genes with significant differential expression. The enrichment factor of an annotation is calculated as the fraction of proteins having this annotation in the sub-network, relative to the entire network. (**b**-**i**) Primary human CD8^+^ T cells were stimulated with CD3/CD28 beads for 3 days in presence or absence of NF-κB specific inhibitor CAPE (**b**, **c**, 5 μM, n = 8), PI3K inhibitor Idelalisib (**d**, **e**, 300 nM, n = 6), Akt inhibitor MK-2206 (**f**, **g**, 200 nM, n = 7), or mTOR inhibitor Rapamycin (**h**, **i,** 600 nM, n = 2). CD8^+^ T cells were stained with antibody against CD8 and TRAIL and were analyzed by flow cytometry. Representative donors are shown in **b**, **d**, **f** and **h**. Data are represented as Mean ± SEM and *p* values were assessed by one-way ANOVA.

We then focused on the involvement of NFκB in HG-enhanced TRAIL expression and tested the effect of the NFκB specific inhibitor CAPE. Notably, the expression of TRAIL on HG-cultured CTLs treated with CAPE was substantially reduced compared to untreated control cells, and even much lower than NG CTLs (Fig. 4b, c). This effect was also confirmed with another NFκB inhibitor MG-132 (Fig. S3). This indicates that NFκB is indispensable for expression of TRAIL in CTLs.

NFκB function can be regulated by the PI3K-Akt pathway, which was among the deregulated KEGG pathways (Spreadsheet 3). We therefore used the PI3Kδ inhibitor idelalisib to block the activity of PI3K. We found that in HG-CTLs, abruption of PI3K function with the corresponding inhibitor reduced the expression of TRAIL to the level of NG or even lower (Fig. 4d, e). We further examined Akt, a molecule downstream of PI3K and upstream of NFκB. We found that disruption of Akt function by MK-2206 abolished HG-enhanced TRAIL expression (Fig. 4f, g). Apart from NFκB, mTOR is also regulated by Akt. To examine whether mTOR is involved in HG-enhanced TRAIL expression, we used rapamycin, a specific inhibitor for mTOR, to functionally block mTOR activity. We found that even with the highest concentration (600 nM), enhancement of TRAIL expression in CTLs by HG was only modestly reduced and remained significantly higher than the level in NG-cultured CTLs (Fig. 4h, i). Taken together, these findings suggest that enhancement of TRAIL expression in CTLs by HG is mediated by the PI3K/Akt/NFκB axis.

### Metformin and vitamin D protect pancreatic beta cells from HG-CTL-mediated apoptosis

Since enhancement of TRAIL levels on CTLs induced by HG leads to apoptosis of pancreatic beta cells as shown above (Fig. 1), a decrease of TRAIL expression to normal levels should protect beta cells. We sought for possible therapeutical approaches to achieve this purpose. We first examined metformin, which is a widely applied first-line medication to treat T2D. We stimulated primary human CD8^+^ T cells in medium containing NG or HG in presence or absence of metformin for three days. We found that TRAIL expression in HG-CTLs was significantly reduced by metformin in a dose-dependent manner (Fig. 5a, b), down to the level of NG-CTLs at 1 mM (compare NG/vehicle and HG/Met 1 mM). These results suggest a putative protective role of metformin on TRAIL-mediated apoptosis of pancreatic beta cells. Apart from metformin, vitamin D has been also linked to diabetes ^25, 26^, glucose metabolism ^27, 28^, protection of pancreatic beta cells ^29^, as well as HG-regulated cell functions ^30^. Therefore, we used 1,25-dihydroxy-vitamin D3 (1,25D3), the active form of vitamin D, to analyze its impact on TRAIL expression. We found that HG-induced enhancement of TRAIL expression on CTLs was reduced by 1,25D3 in a dose-dependent manner (Fig. 5c, d). Notably, the difference in TRAIL expression between HG- and NG-CTLs was completely eliminated by 1,25D3 at 10 µM (Fig. 4c, d, compare NG/vehicle and HG/1,25D3 10 µM). Beside the naturally existing active form 1,25D3, the vitamin D analogue calcipotriol is widely used topically to treat psoriasis. Our results show that nanomolar concentrations of calcipotriol were sufficient to abolish HG-induced enhancement in TRAIL expression (Fig. 5e, f). Our data suggest that activation of the vitamin D pathway can reverse the effect of HG on TRAIL expression on CTLs.

**Figure 5.**
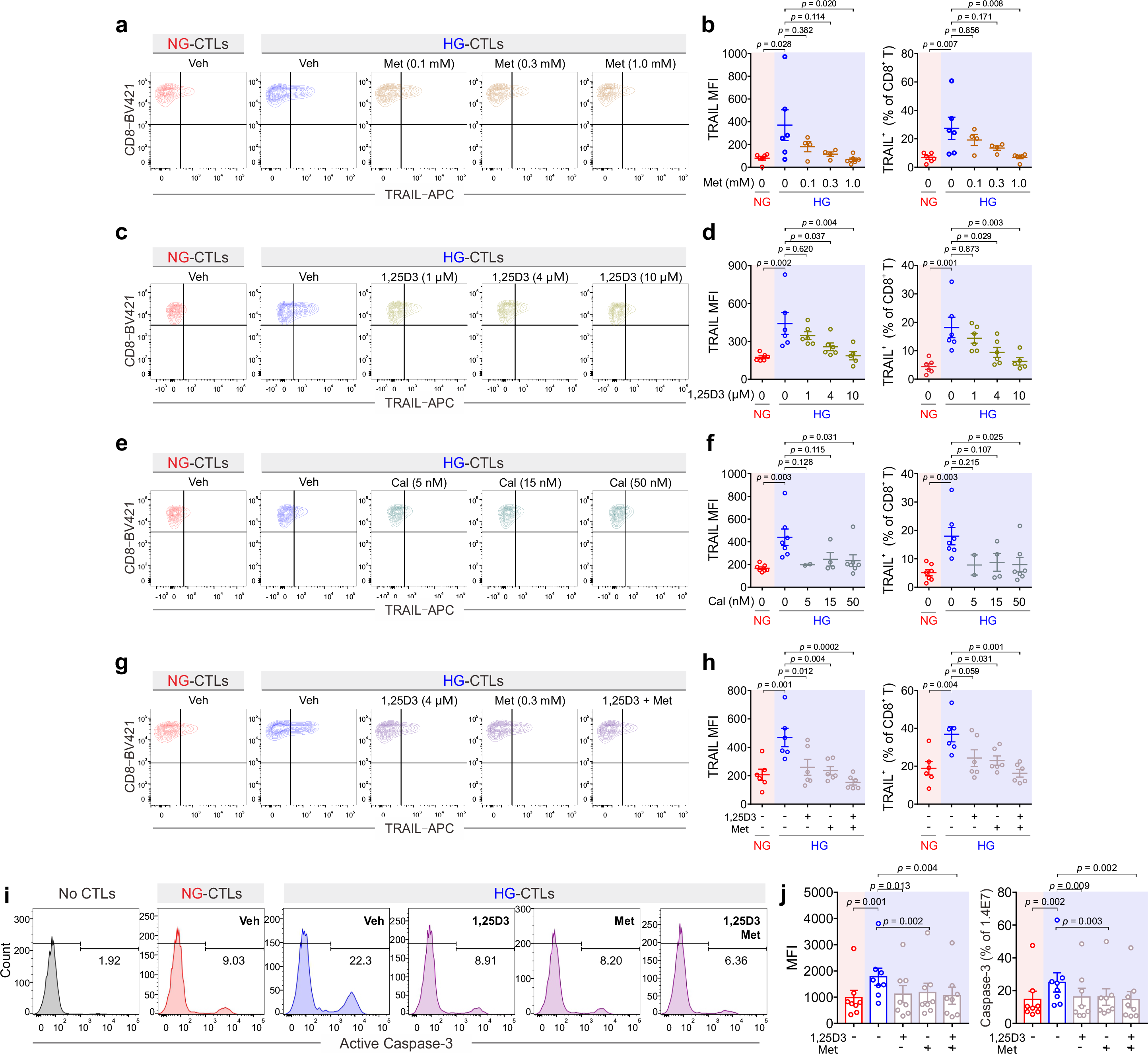
Treatment of metformin and vitamin D rescues TRAIL-mediated pancreatic beta cell destruction. Primary human CD8^+^ T cells were stimulated with CD3/CD28 beads for 3 days in presence of metformin (Met), 1,25D3, or calcipotriol (Cal) with indicated concentrations. TRAIL expression was analyzed by flow cytometry. (**a, b**) Treatment of metformin reduces HG-enhanced TRAIL expression in CD8^+^ T cells (n = 4-6). (**c, d**) Vitamin D regulates TRAIL expression (n = 5-6). (**e**, **f**) Enhancement of TRAIL expression by HG could be downregulated by calcipotriol (n = 2-7). (**g, h**) The combination of Metformin and Vitamin D synergistically inhibits HG-enhanced TRAIL expression (n = 6). (**i, j**) Metformin and Vitamin D can rescue HG-enhanced CTL-mediated beta cell apoptosis. Primary human CD8^+^ T cells were stimulated with CD3/CD28 beads for 3 days in presence of metformin (Met, 300 µM) and/or Vitamin D (1,25D3, 4 µM). 1.4E7 beta cells were co-incubated with CTLs for 4 hours. Results are from 8 donors. Data are shown as means ± SEM. One-way ANOVA was applied for statistical analysis.

Since both metformin and vitamin D down-regulate HG-enhanced TRAIL expression, we tested whether they may function in a synergistic manner. We chose intermediate concentrations for metformin (0.3 mM) and 1,25D3 (4 µM), at which the down-regulation of TRAIL was modest in both cases (Fig. 5b, d). Under this condition, we observed that metformin and 1,25D3 together synergistic down-regulated TRAIL expression enhanced by HG (Fig. 5g, h). Importantly, treatment of metformin or 1,25D3 did not affect T cell activation (Fig. S4a-d) or viability (Fig. S4e). These results demonstrate that metformin and vitamin D function in a synergistic manner to diminish HG-enhanced TRAIL expression on CTLs.

Our findings raise the important question whether metformin and/or vitamin D could protect pancreatic beta cells from HG-cultured CTLs by reducing their TRAIL expression. To address this question, we stimulated primary human CD8^+^ T cells in presence of metformin and/or 1,25D3. Subsequently CTLs were added to 1.4E7 beta cells without metformin and 1,25D3 to avoid the possible effect on beta cells per se. Analysis of 1.4E7 apoptosis at various time points shows that metformin, 1,25D3, or the combination of both reduced TRAIL-mediated beta cell apoptosis to the level of NG control cells (Fig. 5i, j; Fig. S4f, g). In summary, our results clearly indicate that apoptosis of beta cells mediated by HG-induced TRAIL^high^-CTLs can be protected by metformin and/or vitamin D.

### Vitamin D plays an essential role in regulating enhanced TRAIL expression in CD8^+^ T cells from diabetic patients

To test the clinical relevance of metformin and vitamin D protection against TRAIL-induced beta cell apoptosis, we analyzed CTLs from the patients diagnosed with diabetes. We found that after stimulation, the expression of TRAIL on diabetic CTLs was significantly higher compared to that of CTLs from healthy individuals, for both NG and HG conditions (Fig. 6a, b). Remarkably, TRAIL expression of diabetic CTLs under NG condition was already comparable to that of healthy CTLs in HG condition (Fig. 6b). TRAIL expression in diabetic CTLs did not differ between Type 1 and Type 2 diabetes (Fig. S5a, b), or between males and females (Fig. S5c, d), and did not correlate with age (Fig. S5e, f).

**Figure 6.**
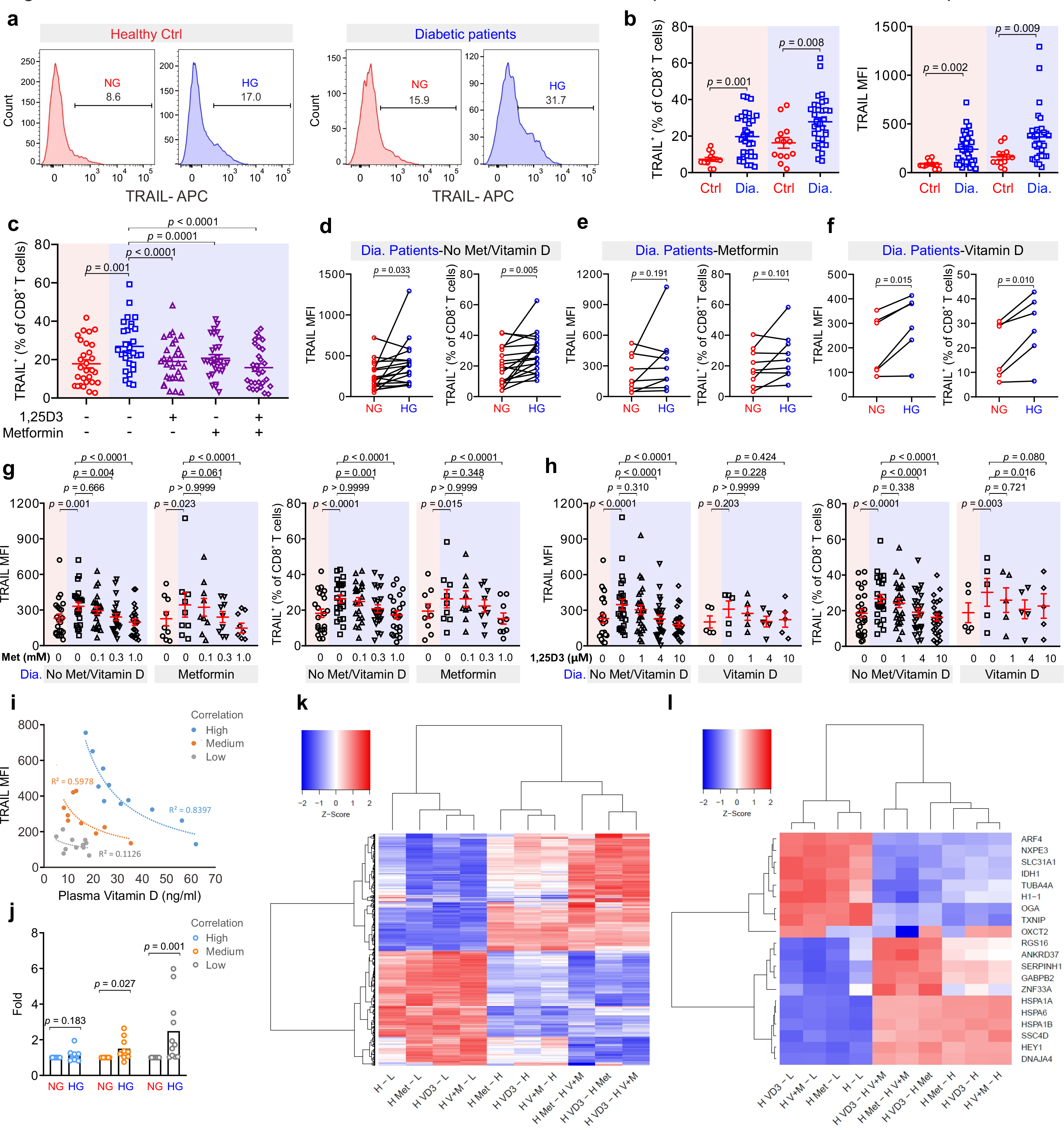
Treatment of metformin and vitamin D decreases TRAIL-expression in CTLs from diabetic patients. (**a, b**) TRAIL expression in CTLs from diabetic patients. PBMCs isolated from healthy individuals or diabetic patients were stimulated with CD3/CD28 beads in NG (5.6 mM, pink shade) or HG (25 mM, blue shade) medium for 3 days. Then the samples were stained with CD3, CD8 and TRAIL and analyzed by flow cytometry (Ctrl, n = 13; diabetic patients, n = 33). One representative donor from either group is shown in **a**. (**c**) Metformin and Vitamin D diminish enhanced TRAIL expression in diabetic CTLs. Diabetic CTLs from **b** were treated with met (300 µM) and/or 1,25D3 (4 µM) during bead-activation for 3 days (n = 29). (**d-h**) Diabetic patients from **b** were categorized into three groups based on whether metformin or vitamin D was taken. (**i, j**) Correlation of TRAIL expression with plasma vitamin D. Mean fluorescence intensity (MFI) of TRAIL in NG-CTLs from diabetic patients shown in **b** is plotted against the concentration of plasma vitamin D of the same patients in **i**. Three groups with different correlation were identified (dashed lines are fitting curves). Change in TRAIL expression by HG in three groups is shown in **j**. Data are represented as Mean ± SEM and *p* values were assessed by two-tailed unpaired Student’s *t* test (**b**), two-tailed paired Student’s *t* test (**d-f, j**), one-way ANOVA (**c**), or two-way ANOVA (**g, h**). R-squared coefficient values were from trendline analysis (**i**). (**k**,**l**) Heatmap of differential gene expression between all pairs of treatments. Listed on the vertical axis are the genes on the microarray. L: NG, H: HG, H+M: HG with metformin, H+D: HG with vitamin D, H+M+D: HG with metformin and vitamin D. Fold changes were standardized by applying a Z-score along the sample axis. Hierarchical clustering (Ward’s method) was carried out for both genes and comparisons (see Spreadsheet 4). The 20 genes with the largest absolute fold change are shown in **l**. Samples were from 7 donors (for L and H) or 3 donors (H+M, H+D, H+M+D).

The correlations between TRAIL expression and diabetes prompted us to examine whether metformin and 1,25D3 can also regulate TRAIL expression in CTLs from diabetic patients. We found that TRAIL expression in diabetic CTLs was down-regulated by 1,25D3 (Fig. S6a), calcipotriol (Fig. S6b) or metformin (Fig. S6c) in a dose-dependent manner. Synergistically, metformin and 1,25D3 could even further suppress TRAIL expression in diabetic CTLs (Fig. 6c). Interestingly, CTLs from patients who did not take metformin or vitamin D showed clear TRAIL enhancement in HG (Fig. 6d), whereas CTLs from patients who took metformin did not show any difference between NG and HG (Fig. 6e). CTLs from patients who took vitamin D exhibited a moderate increase in TRAIL expression under HG condition (Fig. 6f). For both the control group (no metformin and no vitamin D) and the metformin group, culturing CTLs from diabetic patients in presence with metformin in HG further down-regulated TRAIL expression (Fig. 6g). Similarly, for the vitamin D group and the control group, culturing the diabetic CTLs with vitamin D in HG decreased TRAIL expression (Fig. 6h). Collectively, these results demonstrate that in patients with diabetes, HG-enhanced TRAIL expression in CTLs can be efficiently down-regulated by metformin and vitamin D, suggesting the combination of metformin and vitamin D as a promising approach to protect pancreatic beta-cells from TRAIL-mediated CTL killing by HG. In addition, we analyzed the correlation between the concentration of vitamin D in the plasma and the expression of TRAIL on CTLs. We found that based on the correlation, the diabetic CTLs cultured in NG could be categorized into three groups: high, medium, and low correlation (Fig. 6i). Intriguingly, for the group of patients, in which TRAIL expression on CTLs was highly correlated with the concentrations of plasma vitamin D (Correlation High), expression of TRAIL was not further enhanced by HG, whereas for the group Correlation Low, TRAIL expression on CTLs was substantially enhanced by HG condition (Fig. 6j). It indicates that in diabetic patients, sensitivity to vitamin D plays a decisive role in responsiveness of CTLs to HG.

To identify which molecules are targeted by metformin and vitamin D to down-regulate TRAIL in HG-CTLs, we compared transcriptomes of CTLs in five conditions: NG (L), HG (H), HG with metformin (H+M), HG with vitamin D (H+D) or HG with metformin and vitamin D (H+M+D) (Fig. 6k, l), where intermediate concentrations of metformin and vitamin D were used. Remarkably, the analyses of KEGG pathways and enriched GO terms show that OXCT2, an important enzyme in ketone body catabolism, was significantly up-regulated in the group of HG with vitamin D (H+D) and in the group of HG with metformin and vitamin D (H+M+D) compared to control HG group (Spreadsheet 4). Of course, due to the small sample size (3 donors), we could not exclude other genes that could be differentially expressed at the whole transcriptomic level affected by vitamin D and/or metformin. However, OXCT2 stands out in our analysis indicates a rather strong association of vitamin D with ketone body catabolism, suggesting a promising key molecule of vitamin D-regulated response to HG in CTLs.

## Discussion

TRAIL is involved in the development of obesity and diabetes ^11^. In humans, the TRAIL family consists of four membrane receptors, TRAIL-R1, -R2, -R3 and -R4. Among them, TRAIL-R1 and -R2, also known as death receptor 4 (DR4) and DR5, are responsible for inducing TRAIL-mediated cell apoptosis, whereas TRAIL-R3 and -R4 are decoy receptors, which lack the cytoplasmic death domain. Thus, the ratio of death receptors over decoy receptors determines the sensitivity or efficacy of TRAIL-mediated cell apoptosis. TRAIL-R1 and -R2 are highly expressed in many tissues including kidney, heart, adipose tissue, and pancreas; in comparison, decoy receptors TRAIL-R3 and -R4 are not detectable in most of the tissues except for immune function-related organs/tissues (e.g. spleen, bone marrow, and lymph nodes) and pancreas ^31^. Infiltration of immune cells, especially CD8^+^ T cells, is positively correlated with the progression of diabetes and related complications ^11^. Therefore, it is likely that TRAIL^high^- CTLs induced by diabetic conditions not only contribute to destruction of pancreatic beta-cells in an antigen-independent manner but are also involved in attacking TRAIL-R1/-R2^high^ cells in different tissues resulting in undesirable complications. Our findings indicate that harnessing TRAIL expression in CTLs can therefore protect insulin-producing beta cells and can also have a positive effect on diminishing incidence and severity of complications.

In many cell types, glucose transporters (Gluts), especially Glut1, are up-regulated upon activation. For example, expression of Glut1 is elevated upon activation in murine CD4^+^ T cells and macrophages to favor glucose glycolysis over oxidative phosphorylation ^32, 33^. In human T cells, expression of Glut1 is reported to be enhanced in CD3/CD28 bead-stimulated T cells compared to naive T cells ^32^. In comparison, our results how that the expression of Glut1 at the protein level is not altered by HG and even marginally down-regulated at the mRNA level. Our microarray data suggest that the expression level of other Gluts is not influenced by HG, at least at the mRNA level. This indicates that in CTLs, glucose transporters are unlikely the target proteins responsive to HG to mediate HG-induced metabolic reprogramming.

Type I interferons (IFNs, e.g., IFN-α and IFN-β) can induce TRAIL expression on T cells ^34^. However, we could not detect differences in IFN-α and IFN-β expression by microarray analysis between CTLs cultured in NG and HG. This indicates that Type I IFNs are very unlikely to be involved in HG-enhanced TRAIL expression in CTLs. Interestingly, compelling evidence show that IFN-α plays a critical role in initiation of T1D ^35–37^. In a mouse model, blockade of IFN-α signaling at prediabetic stage prevents beta-cells from destruction by CTLs ^38^. Therefore, we postulate that prior to clinical disease-resulted elevation of blood glucose, TRAIL expression on CTLs could be induced by IFN-α, which could also contribute to development of diabetes.

ROS is a byproduct of cellular metabolic redox reaction, which is closely related to progression of diabetes and its complications ^39^. In Th17 cells, high glucose-induced autoimmune reaction is mediated by ROS originated from mitochondria ^40^. Our results show that for human CTLs, ROS is indeed involved in regulating HG-enhanced TRAIL expression, but cytosolic rather than mitochondrial ROS appears to be decisive. This suggests that Th17 cells and CTLs may differ in their regulatory mechanisms in response to HG. Interestingly, ROS can activate the PI3K/Akt pathway as well as the NFκB pathway^41, 42^, which we also found to be essential for HG-enhanced TRAIL expression. Thus, in CTLs, ROS could act upstream of the PI3K/Akt/NFκB axis to regulate HG-enhanced TRAIL expression.

Vitamin D is closely related to insulin resistance and onset of diabetes ^43^. Vitamin D also has a profound impact on immune system ^44^. In particular, expression of vitamin D receptor induced by initial TCR signaling via p38 is essential for upregulation of phospholipase C gamma 1, which plays a pivotal role in T cell activation and Treg/Th17 differentiation ^45, 46^. Interestingly, vitamin D is involved in maintaining mitochondrial functions to optimize cellular redox condition, thus protecting cells from oxidative stress-related damages ^47^. We postulate that vitamin D could diminish HG-enhanced TRAIL expression through reducing intracellular ROS production.

The impact of combined use of vitamin D and metformin has been investigated in various pathological scenarios. For example, combination of vitamin D and metformin shows a positive chemopreventive effect on colorectal cancer in rat and mouse models ^48^ and could be potentially effective to treat patients with polycystic ovary syndrome based on its effect on menstrual cycle ^49^. A recent study reported that in a type 2 diabetes mouse model, which cannot be controlled by metformin alone, additional vitamin D therapy improved insulin sensitivity in skeletal muscles ^50^. This finding supports our findings that combination of metformin and vitamin D could be beneficial to diabetic patients by reducing HG-enhanced expression of TRAIL in CTLs to protect beta cells from TRAIL-mediated apoptosis. In summary, we present a novel mechanism of CTL involvement in progression of diabetes, which establishes CTLs as a target for combined metformin and vitamin D therapy to protect pancreatic beta cells of diabetic patients.

## Acknowledgments

We thank the Institute for Clinical Hemostaseology and Transfusion Medicine for providing donor blood; Carmen Hässig, Cora Hoxha, Gertrud Schäfer and Susanne Renno for excellent technical help. This project was funded by the Deutsche Forschungsgemeinschaft (SFB 1027 A2 to B.Q., A11 to M.H., C3 and ZX to V.H., TRR219 (M04) to L.P.R.), and HOMFOR2019 (to B.Q.). The flow cytometer was funded by DFG (GZ: INST 256/423-1 FUGG). L.S.B. and D.B. are funded by the FNR, respectively by the PRIDE (PRIDE/11012546/NEXTIMMUNE) and the ATTRACT program (A14/BM/7632103).

## Ethical considerations

Research carried out for this study with material from healthy donors (leukocyte reduction system chambers from human blood donors) and diabetic patients is authorized by the local ethic committee (declaration from 16.4.2015 (84/15, Prof. Dr. Rettig-Stürmer) and Ha 84/19, Dr. Christidis, respectively).

## Supplementary Information

**Sup Fig.1.**
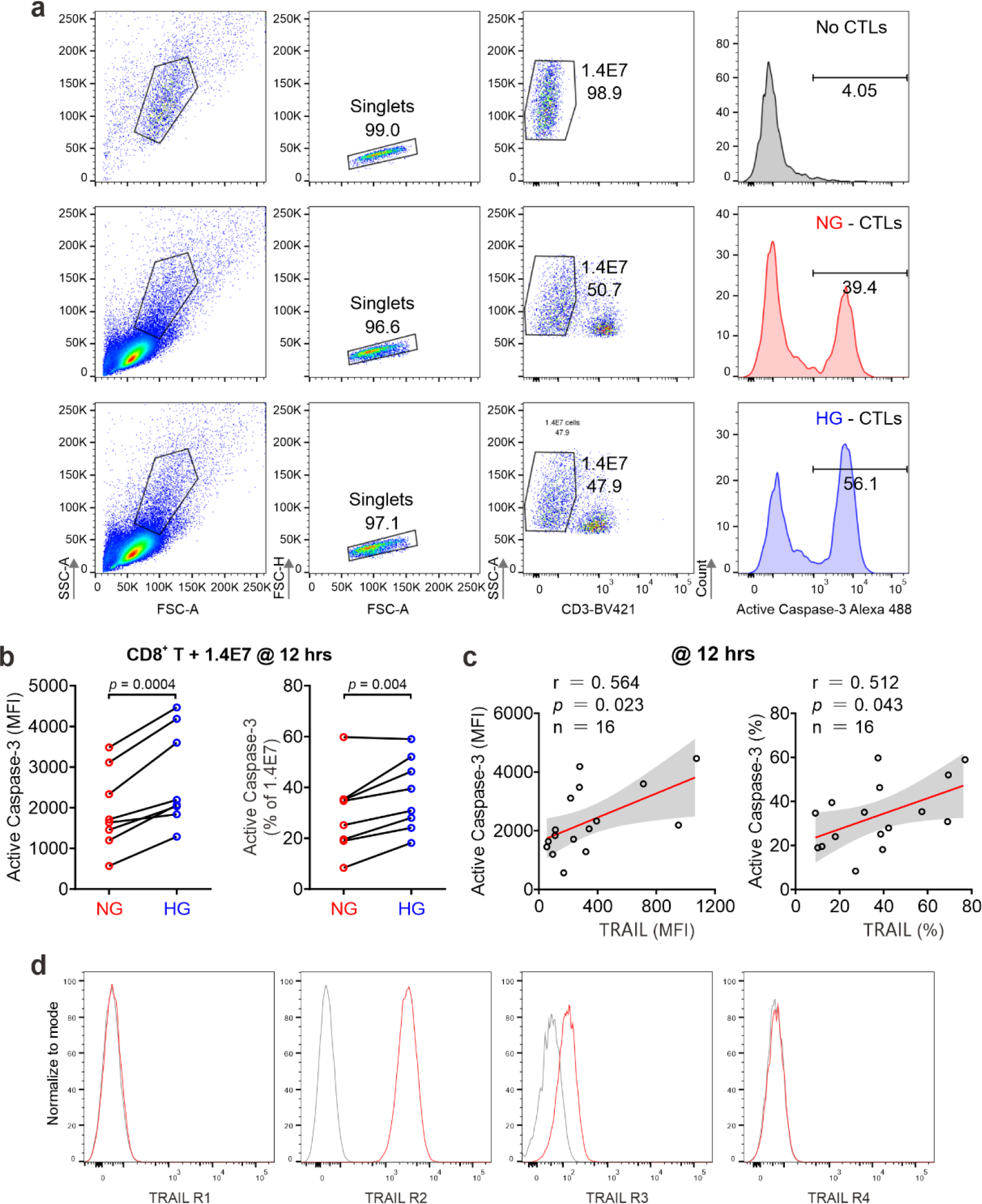
HG-induced TRAIL up-regulation in CD8^+^ T cells enhances beta cell apoptosis. (**a**) Gating for apoptosis assay in 1.4E7 cells at 12 hours. (**b, c**) Apoptosis of pancreatic beta cell induced by CD8^+^ T cells at 12 hours is shown in **c**. The correlation between TRAIL expression in CD8^+^ T cells and 1.4E7 apoptosis of **b** is shown in **c**. Data were analyzed by two-tailed paired Student’s *t* test (**b**), or Pearson’s correlation coefficients (**c**). (**d**) Expression of TRAIL receptors in 1.4E7 beta cells. Gray: IgG control.

**Sup Fig.2.**
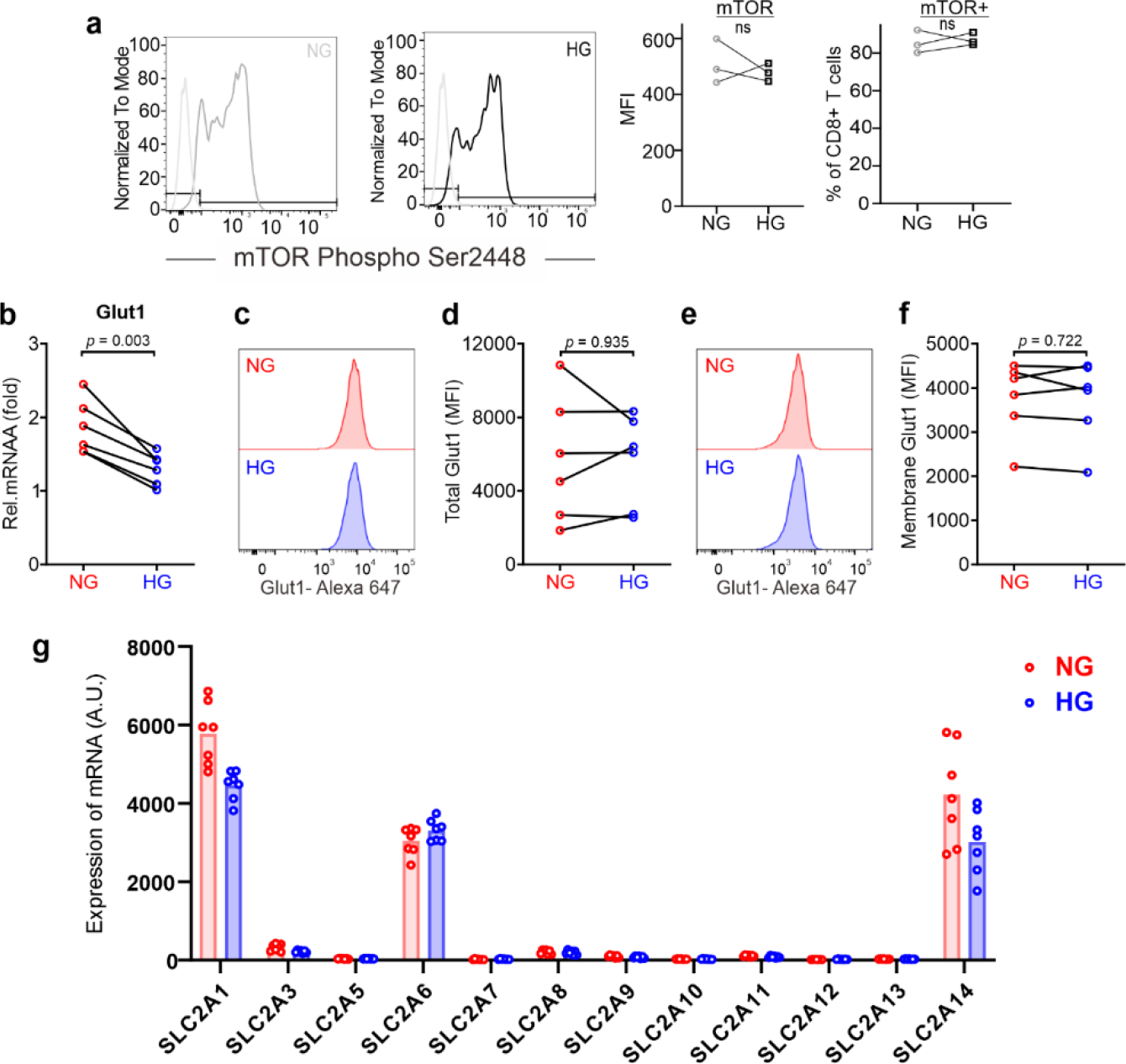
Activation of mTOR and expression of glucose transporters in CTLs. (**a**) Activity of mTOR in CTLs was not altered by HG. PBMCs were stimulated by CD3/CD28 beads and cultured in NG- or HG-medium for 3 days. Activity of mTOR was assessed by mTOR Phospho Ser2448. PBMCs were stained with anti-mTOR Phospho Ser2448, anti-CD3 and anti-CD8 antibody. CD3^+^CD8^+^ cells were gated for analysis. MFI: mean fluorescence intensity. (**b**) Transcriptional levels of Glut1 in CTLs were quantified by qRT-PCR. (**c-f**) Glut1 expression in CTLs in total (**c**, **d**) or on the surface (**e**, **f**) was determined by flow cytometry. Data were analyzed by two-tailed paired Student’s *t* test (**b**, **d**, **f**). (**g**) Expression of glucose transporters at mRNA level. Data are from microarray (Supplementary Table 2). A.U. arbitrary units. Results are from 3 donors (**a**), 6 donors (**b**-**f**), or 7 donors (**g**).

**Sup Fig.3.**
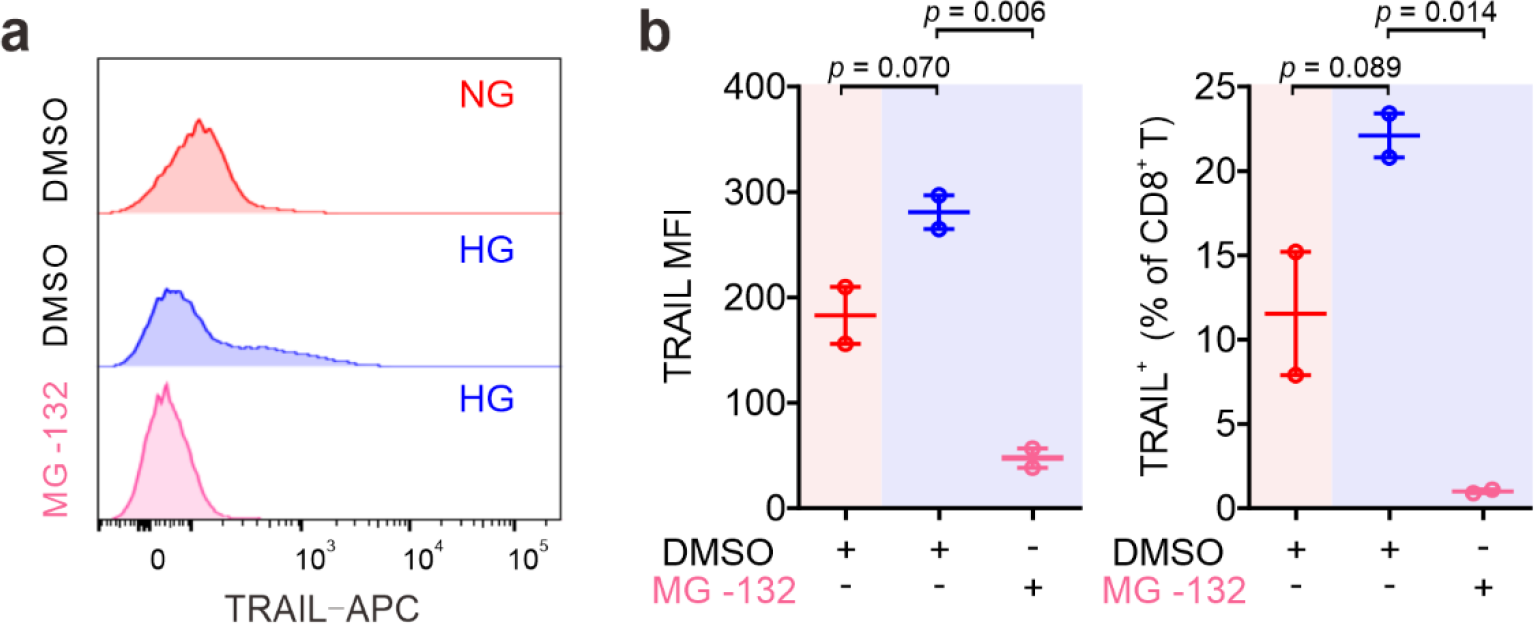
The impact of NKκB inhibitor MG-132 on the TRAIL expression. Primary human CD8^+^ T cells were stimulated with CD3/CD28 beads for 3 days in presence of NKκB inhibitor MG-132 (250 nM). TRAIL expression was analyzed by flow cytometry. One representative donor out of two is shown in **a**. Results are represented as Mean ± SEM. Data were analyzed by one-way ANOVA.

**Sup Fig.4.**
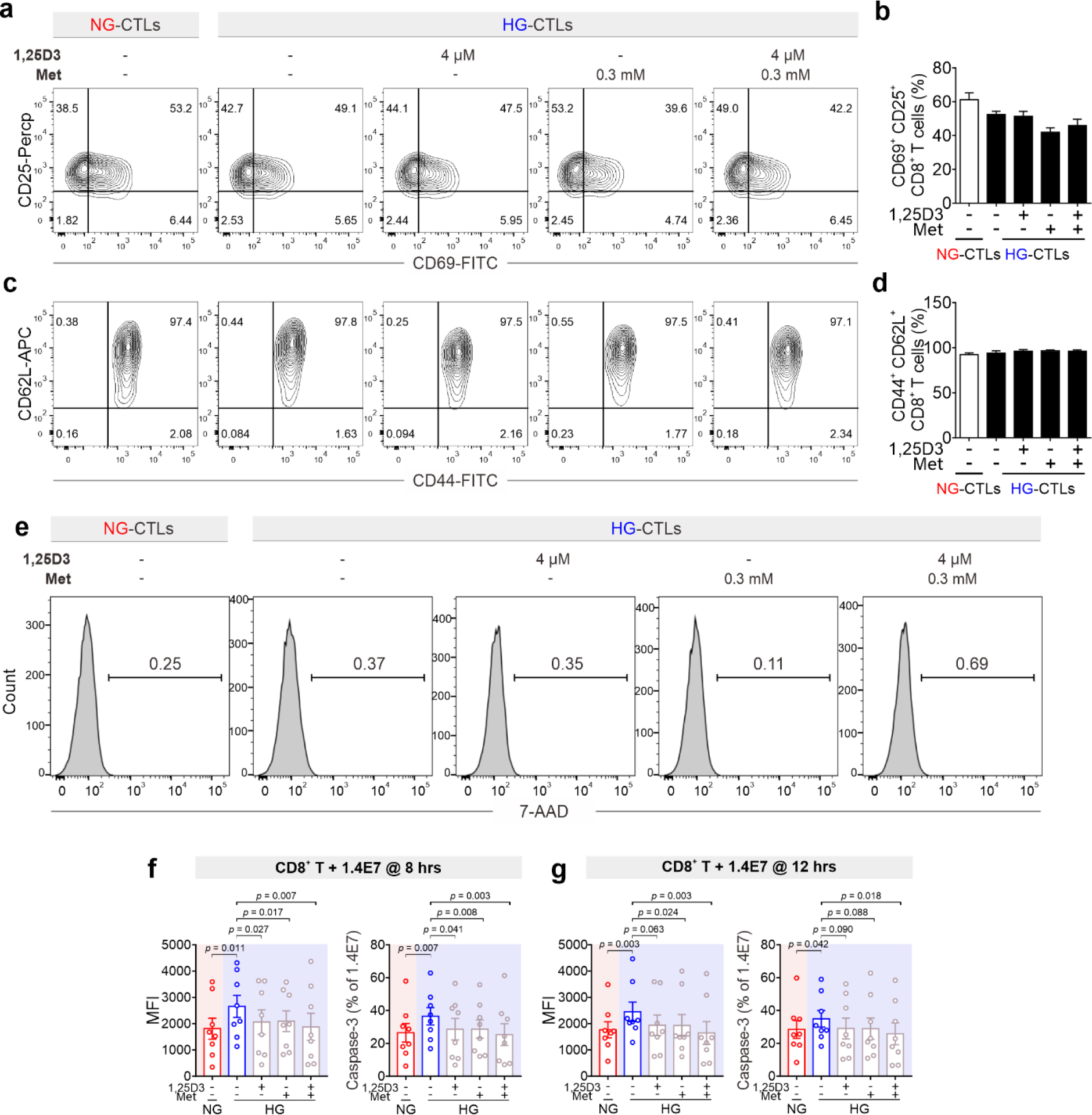
Metformin and vitamin D does not alter activation or viability of CTLs. Primary human CD8^+^ T cells were stimulated with CD3/CD28 beads in NG (5.6 mM) or HG (25 mM) for 3 days in presence with 1,25D3 (4 µM) and/or metformin (300 µM). (**a-d**) Activation of CD8^+^ T cells (n = 3-4 donors). Results are represented as Mean ± SEM. (**e**) Apoptosis of CD8^+^ T cells. (**f**, **g**) Met and 1,25D3 can rescue HG-enhanced beta cell apoptosis by CD8^+^ T cells. Apoptosis of 1.4E7 beta cells was determined by the activity of Caspase-3 at 8 hour (**f**) and 12 hours (**g**) (n = 8 donors). Data were analyzed by one-way ANOVA.

**Sup Fig.5.**
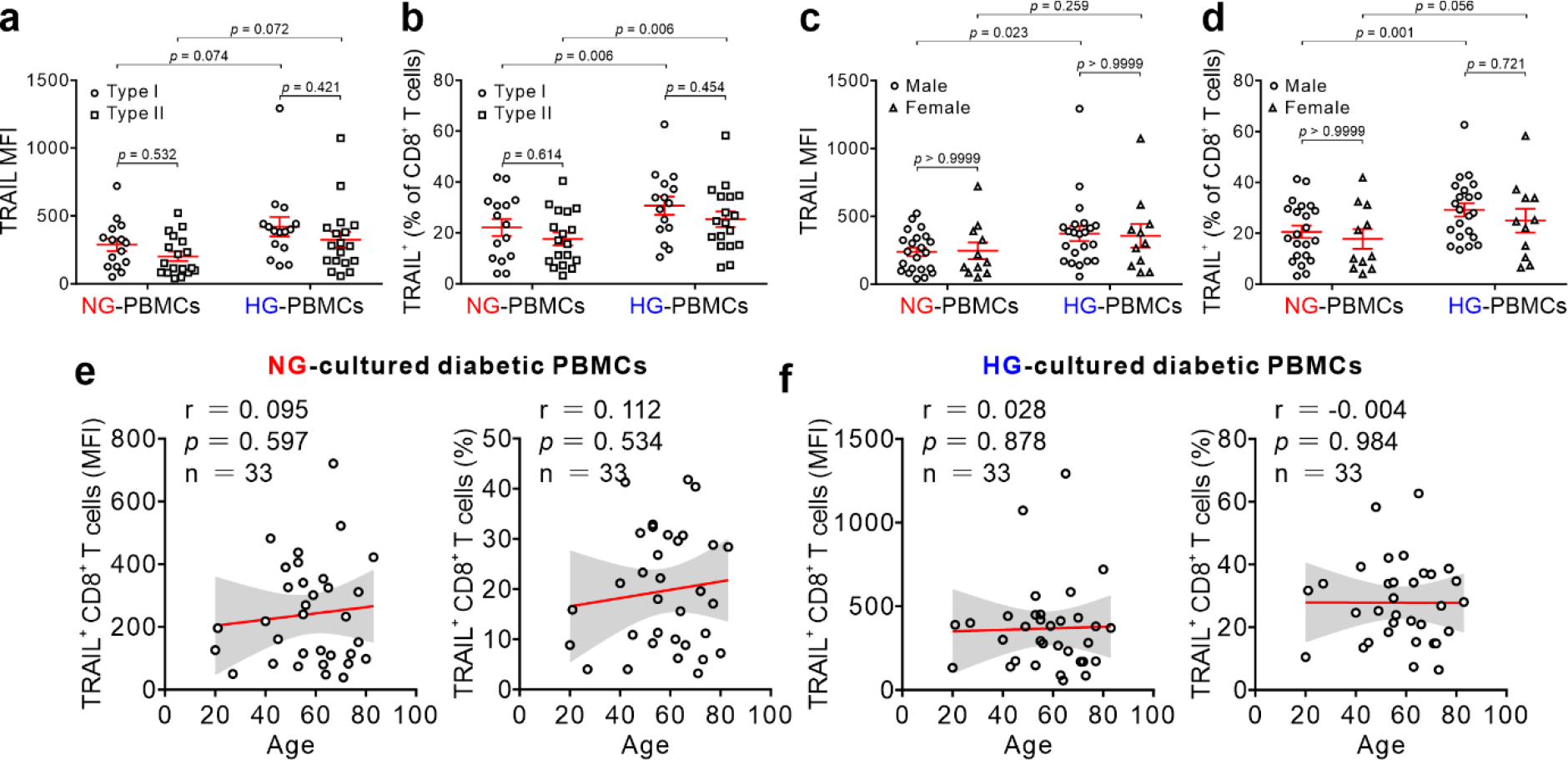
Treatment with met or 1,25D3 can abolish HG-enhanced TRAIL expression in CD8+ T cells of diabetic patients. TRAIL expression is not correlated with types (**a**, **b**), gender (**c**, **d**) or age (**e**, **f**). PBMCs from diabetic patients same as in Fig. 5b. Results were represented as Mean ± SEM. Data were analyzed by two-way ANOVA (**a**-**d**), or Pearson’s correlation coefficients (**e**, **f**).

**Sup Fig.6.**
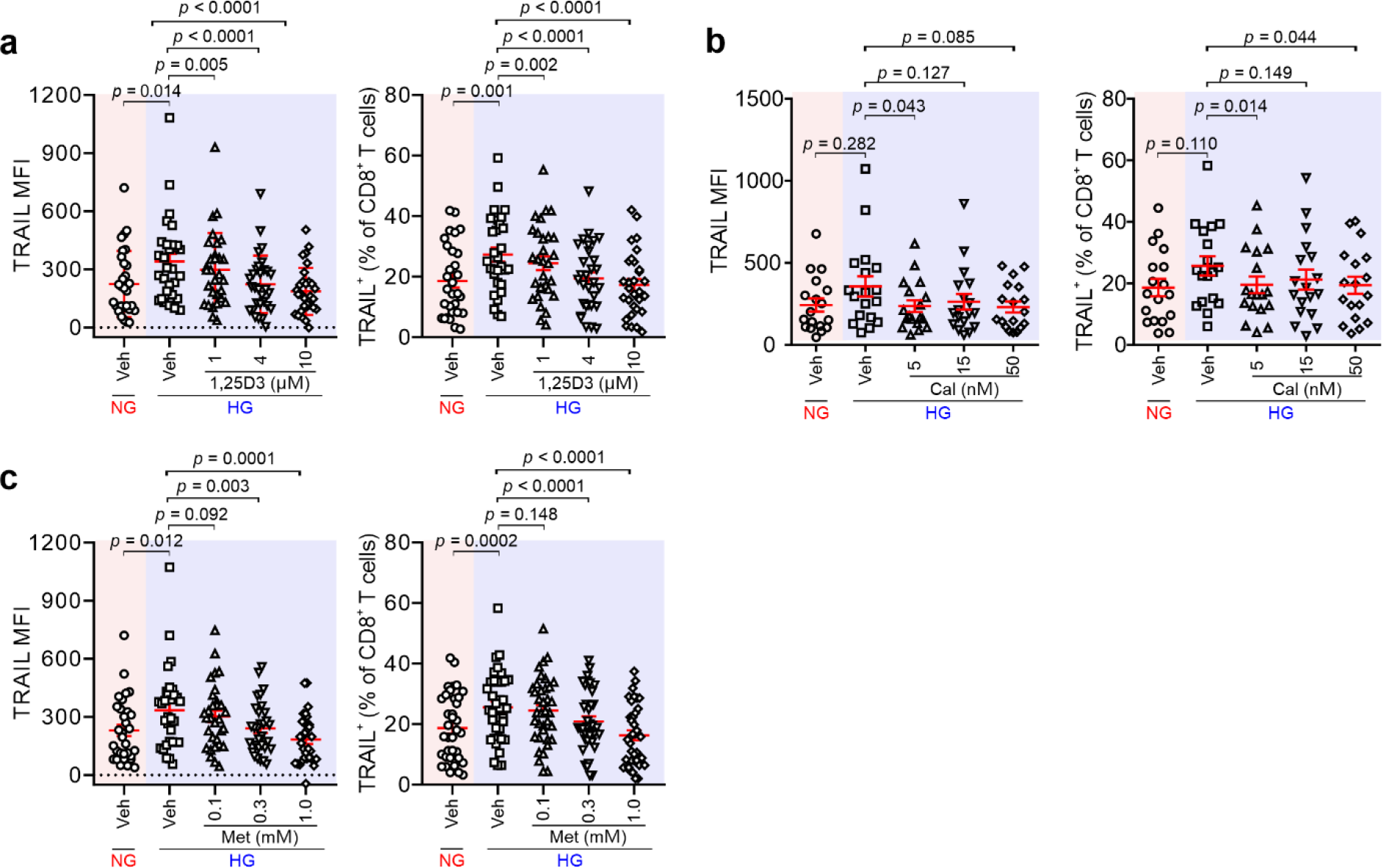
Impact of Metformin or vitamin D pathway on HG-enhanced TRAIL expression in CD8^+^ T cells. PBMCs from diabetic patients were stimulated with CD3/CD28 beads in presence of vitamin D (1,25D3, **a**), calcipotriol (Cal, **b**) metformin (Met, **c**) for 3 days. TRAIL expression was determined by flow cytometry. Results are represented as Mean ± SEM (1,25D3 n = 11; Cal n = 18; Met n = 11). Data were analyzed by one-way ANOVA.

## Spreadsheet legends

**Spreadsheet 1. Normalized gene expression data.** Microarray gene expression data were processed with NormExp background correction and quantile normalization. The control probes were removed, as well as probes that were not expressed in at least three of 23 samples. Remaining probes were annotated with gene symbols and entrez identifiers, according to the specifications of the microarray platform (Agilent design number 026652).

**Spreadsheet 2. GO term enrichment analysis results.** Enriched Gene Ontology (GO) terms for pairwise comparisons between conditions. GO term enrichment analysis was performed with the goana function from the limma R-package. Each term was annotated with its respective up- and down-regulated genes.

**Spreadsheet 3. KEGG pathway enrichment analysis results.** Enriched KEGG pathways for each pairwise comparison between conditions. Pathway enrichment analysis was performed with the kegga function from the limma R-package. Each pathway was annotated with its respective up- and down-regulated genes.

**Spreadsheet 4. Differential gene expression analysis results.** Results of pairwise differential expression analysis. Contains the average log2-intensity for each gene (A), along with the log2 fold change coefficients for all treatment pairs (Coef), the moderated t-statistics (t), the unadjusted p-values (p-value), the p-values after Benjamini-Hochberg correction (p.value.adj), the F statistic results (F, F.p.value), and a classification for each pairwise treatment comparison into up-regulated (1), down-regulated (-1) and unregulated (0). Additionally, the probes were annotated with gene symbols and entrez identifiers. The analysis was carried out with the contrasts.fit, eBayes and decideTests functions from the limma R-package.

